# Transmission dynamics of MERS-CoV in a transgenic human DPP4 mouse model

**DOI:** 10.1101/2023.11.22.568286

**Authors:** Neeltje van Doremalen, Trenton Bushmaker, Robert J. Fischer, Atsushi Okumura, Dania Figueroa, Rebekah J. McMinn, Michael Letko, Greg Saturday, Vincent J. Munster

## Abstract

Since 2002, three novel coronavirus outbreaks have occurred: severe acute respiratory syndrome coronavirus (SARS-CoV-1), Middle East respiratory syndrome coronavirus (MERS-CoV), and SARS-CoV-2. A better understanding of the transmission potential of coronaviruses will result in adequate infection control precautions and an early halt of transmission within the human population. Experiments on the stability of coronaviruses in the environment, as well as transmission models, are thus pertinent. Here, we show that transgenic mice expressing human DPP4 can be infected with MERS-CoV via the aerosol route. Exposure to 5×10^6^ TCID_50_ and 5×10^4^ TCID_50_ MERS-CoV per cage via fomites resulted in transmission in 15 out of 20 and 11 out of 18 animals, respectively. Exposure of sentinel mice to donor mice one day post inoculation with 10^5^ TCID_50_ MERS-CoV resulted in transmission in 1 out of 38 mice via direct contact and 4 out of 54 mice via airborne contact. Exposure to donor mice inoculated with 10^4^ TCID_50_ MERS-CoV resulted in transmission in 0 out of 20 pairs via direct contact and 0 out of 5 pairs via the airborne route. Our model shows limited transmission of MERS-CoV via the fomite, direct contact, and airborne routes. The hDPP4 mouse model will allow assessment of the ongoing evolution of MERS-CoV in the context of acquiring enhanced human-to-human transmission kinetics and will inform the development of other transmission models.

## Introduction

In the last two decades, three novel coronaviruses have caused outbreaks in the human population. Severe acute respiratory syndrome coronavirus (SARS-CoV-1) was first identified in 2003 after it caused human cases in the Guangdong province, China, in November 2002. From Guangdong, the virus spread to 37 countries, resulting in >8000 infected people with a case fatality rate of 9.5% ^1^. Middle East respiratory syndrome coronavirus (MERS-CoV) was first detected in 2012 and still circulates in the dromedary camel population, from which it infects the human population. Since 2012, more than 2600 cases with a case fatality rate of 36% ^2^ have been reported. The largest pandemic with a coronavirus to date started in December 2019 and was caused by SARS-CoV-2. To date, more than 770 million cases have been reported, resulting in nearly 7 million deaths ^3^.

A better understanding of the transmission potential of coronavirus is crucial when devising personal protection equipment for healthcare staff and quarantine measurements. The stability of MERS-CoV, SARS-CoV-1, and SARS-CoV-2 has been investigated under several different environmental conditions in both fomites and aerosols ^4–6^. Experimental transmission models have been developed for SARS-CoV-2 and focus mainly on hamsters and ferrets ^7–13^, whereas SARS-CoV-1 transmission has been shown in ferrets and cats ^13,14^. However, transmission of MERS-CoV in animal models has not yet been reported. A review of 681 MERS cases in the Kingdom of Saudi Arabia (KSA) estimated that 12% of cases were infected via direct exposure to dromedary camels, and 88% resulted from human-to-human transmission ^15^. Analysis of transmission dynamics showed that the number of subsequent generations is limited. The risk of a human-to-human transmission event differs per generation: the initial zoonotic transmission risk is 22.7%. Then it drops to 10.5% for the second generation, 6.1% for the third generation, and 3.9% for the fourth generation ^16^. This shows that although human-to-human transmission contributes significantly to the number of human MERS-CoV cases, transmission is not sustained. Human-to-human transmission of MERS-CoV can be divided into household-associated healthcare-associated (nosocomial transmission). Epidemiological modeling of MERS-CoV transmission estimates nosocomial transmission to be ten times higher than household transmission ^17^.

Virus transmission can occur via different routes, including fomites, direct contact, and aerosols. Knowledge on the transmission routes of emerging coronaviruses is essential in designing broad preemptive countermeasures against zoonotic and human-to-human transmission. More specifically, it will improve the ability of hospitals to reduce the likelihood of human-to-human transmission by implementing appropriate personal protective equipment and hospital hygiene procedures.

The best transmission models for SARS-CoV-2 are the hamster and ferret models ^7–13^. However, these animals are not naturally susceptible to MERS-CoV ^18,19^ and likely will require expression of hDPP4. Although transmission in mice is likely more limited than in hamsters or ferrets, SARS-CoV-2 transmission has been shown in mice ^20,21^. Therefore, we utilize a transgenic mouse model expressing human DPP4 (hDPP4 mice) to investigate whether MERS-CoV can transmit via fomites, direct contact, or the airborne route.

## Materials and Methods

### Ethics statement

Approval of animal experiments was obtained from the Institutional Animal Care and Use Committee of the Rocky Mountain Laboratories. Performance of experiments was done following the guidelines and basic principles in the United States Public Health Service Policy on Humane Care and Use of Laboratory Animals and the Guide for the Care and Use of Laboratory Animals. Work with infectious MERS-CoV strains under BSL3 conditions was approved by the Institutional Biosafety Committee (IBC). Inactivation and removal of samples from high containment was performed per IBC-approved standard operating procedures.

### Development of hDPP4 mice

A ROSA26 knock-in vector (ingenious Targeting Laboratory) containing a 3’ splice acceptor, *LoxP*-flanked neomycin stop cassette, Kozak sequence, human DPP4 cDNA sequence and bovine growth hormone poly-A tail was injected into balb/c embryonic stem (ES) cells via electroporation. ES cells were injected into balb/c blastocysts. Resulting chimeric mice were bred with wildtype balb/c mice. Heterozygous offspring were bred with BALB/c-Tg(CMV-cre)1Cgn/J mice (Jackson Laboratory) to produce mice ubiquitously expressing hDPP4. Deletion of the *LoxP*-flanked neomycin stop cassette occurs in all tissues, including germ cells, in BALB/c-Tg(CMV-cre)1Cgn/J mice. Polymerase chain reaction (PCR) was performed to genotype each mouse using a three primer set-up; forward primer (FP) AGCACTTGCTCTCCCAAAGTC, reverse primer 1 (RP1) GACAACGCCCACACACCAGGTTAG and reverse primer 2 (RP2) TCTTCTGTAATCAGCTGCCTTTTA.

### Virus and cells

HCoV-EMC/2012 was kindly provided by Erasmus Medical Center, Rotterdam, The Netherlands. Virus propagation was performed in VeroE6 cells in DMEM (Sigma) supplemented with 2% fetal calf serum (Logan), 1 mM L-glutamine (Lonza), 50 U/ml penicillin and 50 μg/ml streptomycin (Gibco) (2% DMEM). VeroE6 cells were maintained in DMEM supplemented with 10% fetal calf serum, 1 mM L glutamine, 50 U/ml penicillin and 50 μg/ml streptomycin. Virus was titrated by inoculating VeroE6 cells with tenfold serial dilutions of virus in 2% DMEM. Five days after inoculation, cytopathic effect (CPE) was scored and TCID_50_ was calculated from four replicates by the method of Spearman-Karber.

### DPP4 expression

The expression of DPP4 in mouse tissue was examined using an enzyme-linked immunosorbent assay (ELISA). Protein was extracted from tissue membranes using the MEM-PER protein extraction kit (89842, ThermoFisher Scientific). Protein extracts were measured using a DPP4 ELISA kit (DY1180 and DY008, R&D resources). Protein concentrations were normalized using the BCA protein assay kit (23225, Pierce).

### Inoculation experiments

hDPP4 mice were inoculated intranasally with MERS-CoV isolate HCoV-EMC/2012 in a total volume of 30 µl. Aerosol inoculation using the AeroMP aerosol management platform (Biaera technologies, USA) was performed as described previously ^18^. Briefly, mice were exposed to a single 10-minute aerosol exposure whilst contained in a stainless-steel wire mesh cage (5 mice per cage, 2 cages per run, mo anesthesia). Aerosol particles were generated by a 3-jet collison nebulizer (Biaera technologies, USA) and ranged from 1-5 µm in size. Respiratory minute volume rates of the animals were determined using Alexander *et al*. ^22^. Weights of the animals were averaged and the estimate inhaled dose was calculated using the simplified formula D = R × C_aero_ × T_exp_ ^23^, where D is the inhaled dose, R is the respiratory minute volume (L/min), C_aero_ is the aerosol concentration (TCID_50_/L), and T_exp_ is duration of the exposure (min). After inoculation, animals were observed daily for signs of disease. Euthanasia was indicated at >20% loss of initial body weight, if severe respiratory distress was observed, or if neurological signs were observed. Oropharyngeal and nasal swabs were collected daily in 1 ml DMEM with 100 U/ml penicillin and 100 µg/ml streptomycin.

### Transmission experiments

Fomite transmission was examined by contaminating cages containing two metal and two plastic discs [3] with 0.5 mL of MERS-CoV isolate HCoV-EMC/2012 (total dose: 5 × 10^6^ or 5 × 10^4^ TCID_50_ per cage) in the following manner: 50 µl of virus was placed on the water bottle, 50 µl of virus was placed on the food, 50 µl of virus was placed per disc, and 4 × 50 µl of virus was placed on the flooring of the cage. Mice were placed in the cages 10 minutes post contamination and followed as described above. Contact and airborne transmission were examined by intranasal inoculation of donors with 10^4^ or 10^5^ TCID_50_ of MERS-CoV. At 1 dpi, sentinel animals were placed in the same cage (contact transmission) or in the same cage on the other side of a divider (airborne transmission) (*Figure S1*). This divider prevented direct contact between the donor and sentinel mouse and the movement of bedding material. Only one transmission pair was housed per cage. Hereafter, mice were followed as described above.

### Histopathology and immunohistochemistry

Murine tissues were evaluated for pathology and the presence of viral antigen as described previously ^24^. Briefly, tissues were fixed in 10% neutral-buffered formalin for 7 days and paraffin-embedded. Tissue sections were stained with hematoxylin and eosin (H&E). An in-house produced rabbit polyclonal antiserum against HCoV-EMC/2012 (1:1000) was used as a primary antibody for the detection of viral antigen. A commercial antibody (AF1180, R&D resources) was used for the detection of DPP4.

### Viral RNA detection

Tissues were homogenized in RLT buffer and RNA was extracted using the RNeasy method on the QIAxtractor (Qiagen) according to the manufacturer’s instructions. RNA was extracted from swab samples using the QiaAmp Viral RNA kit on the QIAxtractor. For one-step real-time qPCR, five μl RNA was used in the Rotor-GeneTM probe kit (Qiagen) according to instructions of the manufacturer. Standard dilutions of a virus stock with known titer were run in parallel in each run, to calculate TCID_50_ equivalents in the samples. Initial detection of viral RNA was targeted upstream of the envelope gene sequence (UpE) ^25^, confirmation was targeted at the ORF1A ^26^. MERS-CoV M-gene mRNA was detected as described by Coleman *et al* ^27^. Values with Ct-values >38 were excluded. For digital droplet PCR (ddPCR, Biorad), eight µl RNA was added to supermix for probes and assay was run according to instructions of the manufacturer using UpE primers and probe allowing absolute quantification of target RNA.

### Serology

Enzyme-linked immunosorbent assay (ELISA) was performed as described previously ^28^. Briefly, spike S1 antigen (Sino Biological Inc., 0.5 µg/mL in 50 mM bicarb binding buffer (4.41 g KHCO_3_ and 0.75 g Na_2_CO_3_ in 1 L water) was bound to MaxiSorb plates (Nunc) and then blocked with 5% non-fat dried milk in PBS-0.1% Tween (5MPT). Serum samples were diluted in 5MPT. Detection of MERS-specific antibodies was performed with HRP-conjugated IgG (H+L) secondary antibody and developing solution (KPL) followed by measurement at 405 nm. Sera was termed seropositive if the optical density value was higher than the average + 3x standard deviation of sera obtained from randomly selected mice before MERS-CoV inoculation.

### Statistical analysis

Statistical analysis was performed by Log-rank (Mantel-Cox) test to compare survival curves, and by Kruskall-Wallis test followed by Mann-Whitney test. P-values<0.05 were considered significant.

## Results

### Intranasal or aerosol inoculation of hDPP4 mice with MERS-CoV

In our hDPP4 mouse model, expression of hDPP4 can be found throughout the respiratory tract and the kidney (*Figure 1a-d*). Inoculation with 10^5^ TCID_50_ MERS-CoV strain HCoV-EMC/2012 resulted in >20% weight loss and 100% lethality (*Figure 1e-f*). Viral gRNA could be detected in oropharyngeal swabs on all six days during the experiment (*Figure 1g*). Infectious virus was detected in lung tissue at three days post-inoculation (dpi) and at endpoint, as well as in brain tissue at end point, but not in the kidney (*Figure 1h*). We then investigated whether inoculation via aerosols would result in productive infection of hDPP4 mice. We inoculated ten mice via aerosols with an estimated 3.8 × 10^2^ TCID_50_ MERS-CoV per mouse. All mice reached the endpoint criteria (*Figure 2a*). Shedding, as measured by gRNA presence in oropharyngeal swabs, could be detected on all days (*Figure 2b*). Viral RNA was detected in lung tissue (four out of four) and brain tissue (one out of four) on three dpi and all tissue on the day of endpoint criteria (*Figure 2c*). Using immunohistochemistry, we compared the cellular tropism of MERS-CoV replication in hDPP4 mice between intranasal (inoculation dose = 10^3^ TCID_50_) and aerosol inoculation (3.8 × 10^2^ TCID_50_) in the upper and lower respiratory tract at three dpi. For both groups, viral replication was detected in lung tissue: type I and type II pneumocytes were positive for MERS-CoV antigen. However, even though we showed that cells lining the trachea, as well as the nasal turbinates, of hDPP4 mice express hDPP4 (*Figure 1a-c*), MERS-CoV antigen staining was negative in the trachea. No staining in the nasal turbinates was detected for the aerosol-inoculated group, whereas staining in the nasal turbinates of the intranasally inoculated group was very limited (*Figure 3*). This MERS-CoV respiratory tropism is similar to what has been observed in humans ^29^; MERS-CoV predominantly targets the cells in the lower respiratory tract.

**Figure 1.**
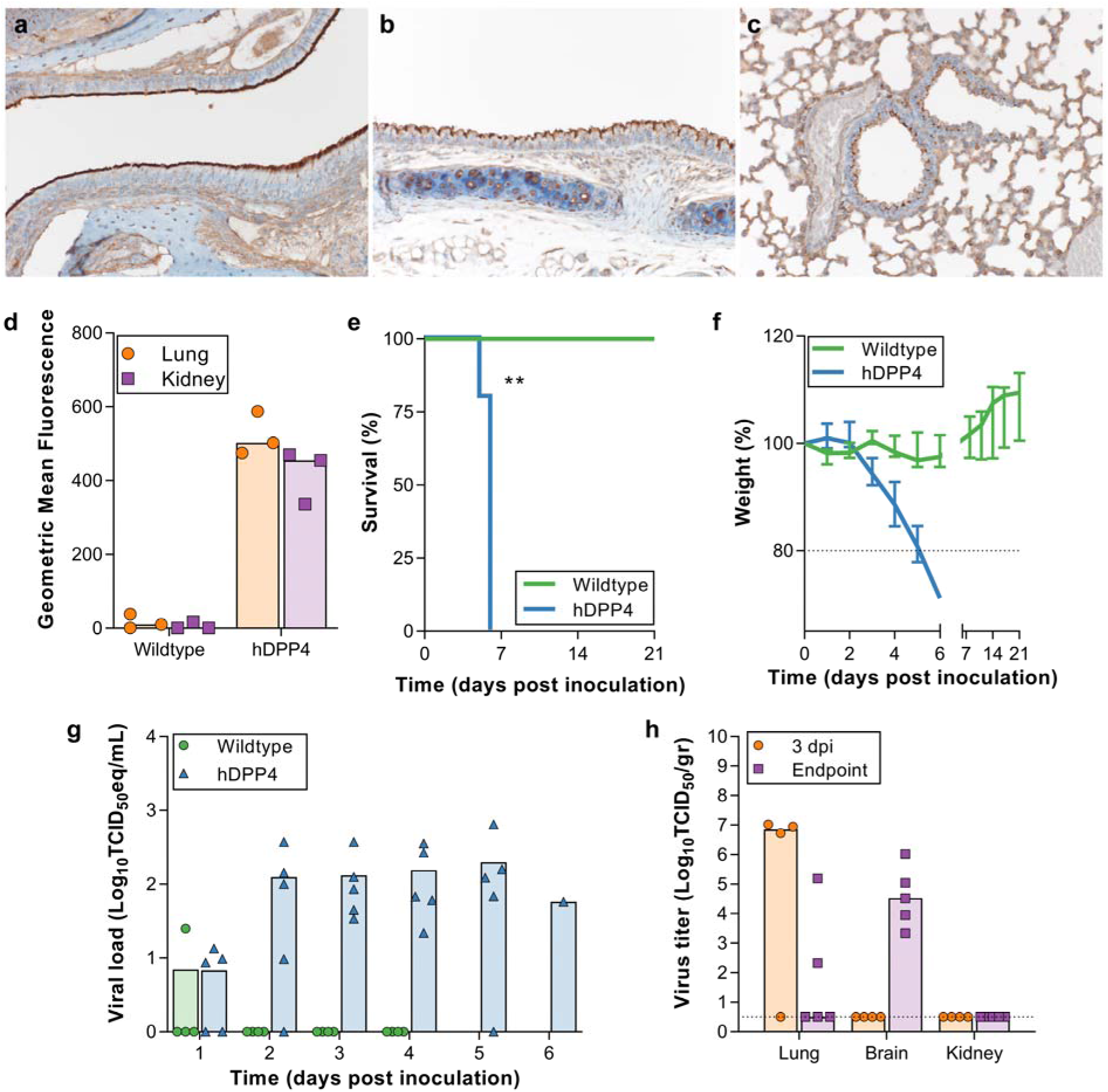
Inoculation of hDPP4 mice, but not wildtype mice, with MERS-CoV results in disease and viral shedding. (a-c) Detection of hDPP4 expression in hDPP4 mice using immunohistochemistry in (a) nasal mucosa; (b) trachea; and (c) type I and II pneumocytes, bronchiolar and endothelial cells in lung tissue. (d) Comparison of hDPP4 expression in lung and kidney tissue obtained from wildtype and hDPP4 mice using flow cytometry. N=3, bars represent median. (e) Survival curves of mice after inoculation with 5×10^5^ TCID_50_ MERS-CoV. Mice were euthanized upon reaching >20% of body weight loss (dotted line). N=4 (Wildtype) or 5 (hDPP4). (f) Relative weight loss in mice after MERS-CoV inoculation. The lines represent median±range. (g) Viral load (gRNA) in oropharyngeal swabs obtained from mice after inoculation with 5×10^5^ TCID_50_ MERS-CoV. (h) Infectious MERS-CoV titers in lung, brain, and kidney tissue of hDPP4 mice. Dotted line = detection limit.

**Figure 2.**
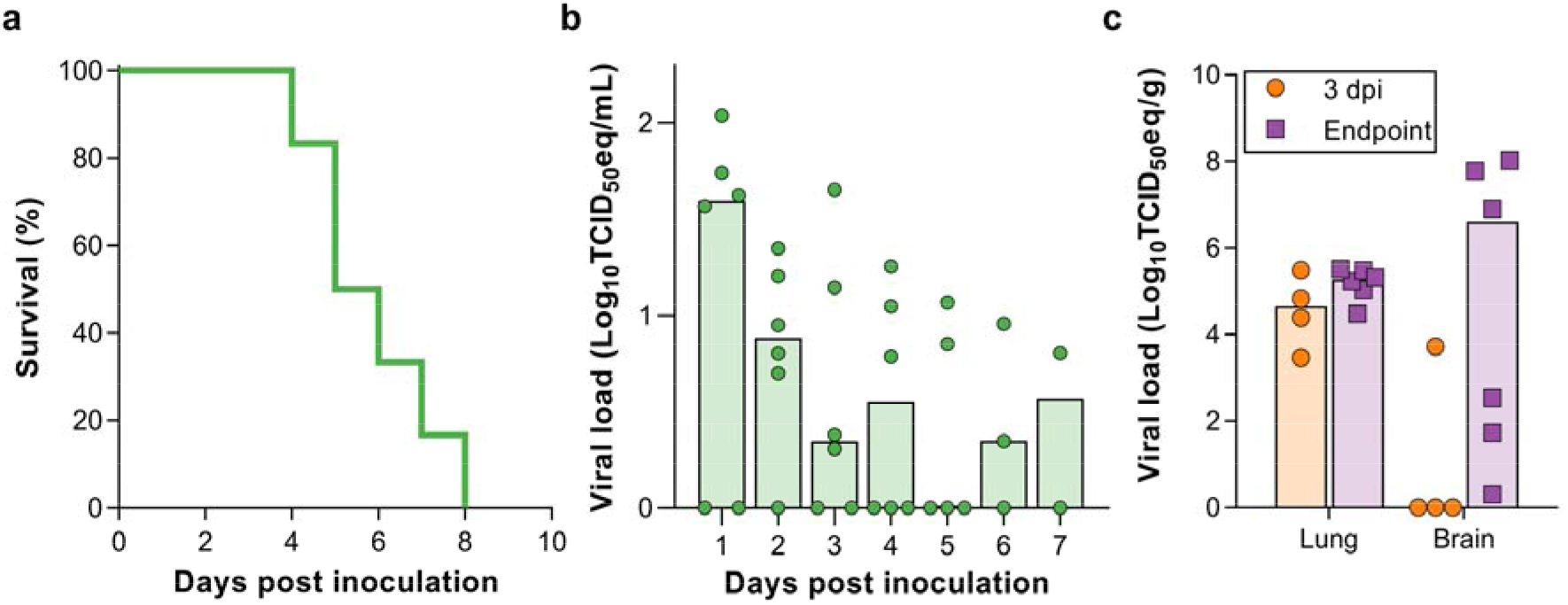
MERS-CoV-associated disease upon inoculation via aerosols. (a) Survival curves of mice inoculated with 3.8 × 10^2^ TCID_50_ MERS-CoV via aerosols. N=6. (b) Viral load (gRNA) in oropharyngeal swabs obtained from mice inoculated with 3.8 × 10^2^ TCID_50_ MERS-CoV via aerosols. Bars represent median. (b) Viral load (gRNA) in lung and brain tissue of hDPP4 mice inoculated with 3.8 × 10^2^ TCID_50_ MERS-CoV via aerosols.

**Figure 3.**
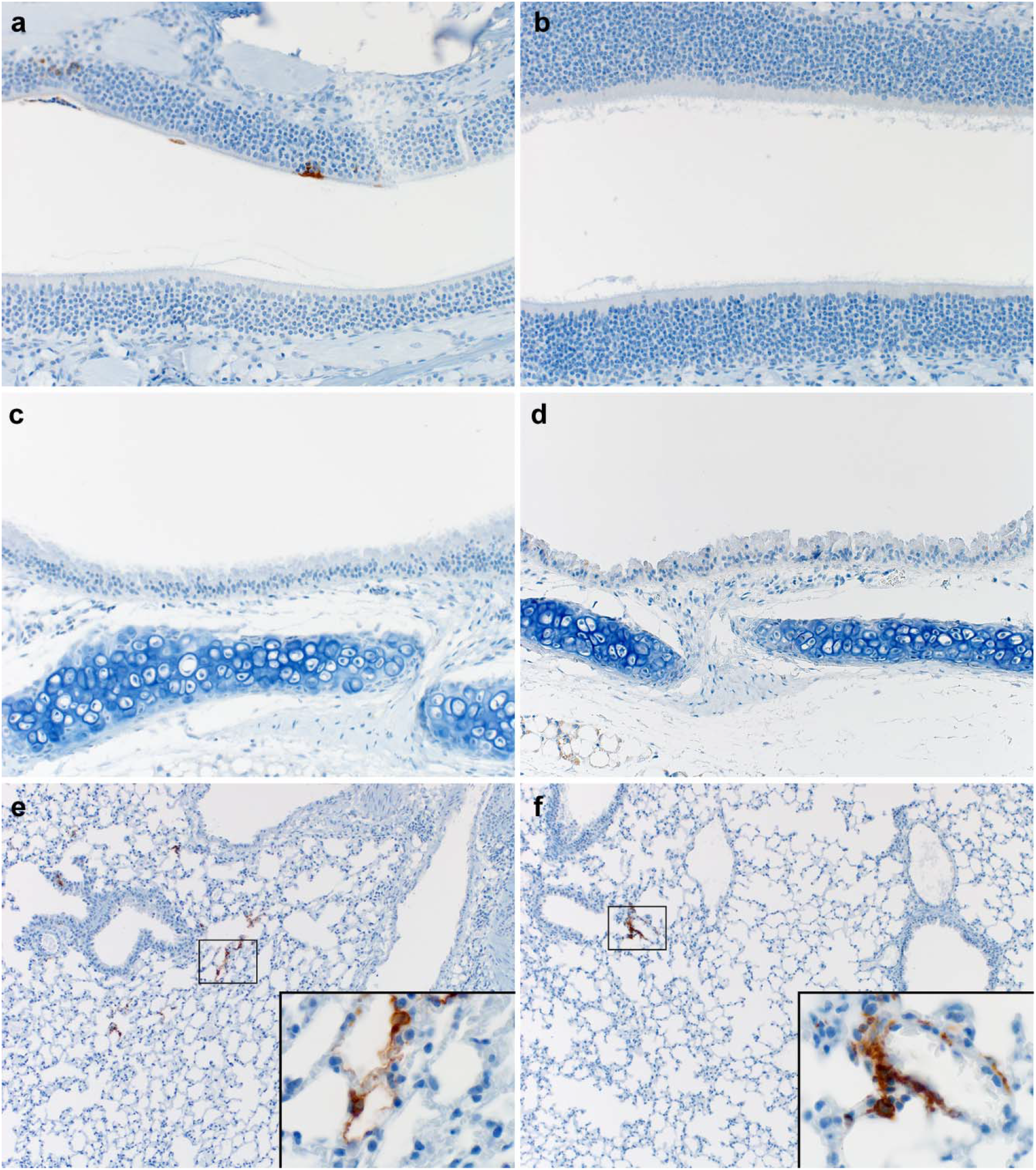
Immunohistochemistry for MERS-CoV antigen in respiratory tract of mice inoculated via the intranasal route or via aerosols. (a-b) Nasal turbinates; (c-d) Trachea; (e-f) Lungs; (a,c,e) Intranasal inoculation; (b,d,f) Aerosol inoculation. Tissues were stained with an in-house produced rabbit polyclonal antiserum against HCoV-EMC/2012 for the detection of viral antigen. Immunohistochemistry staining reveals MERS-CoV antigen in type I and type II pneumocytes and limited staining in nasal turbinates (image represents only staining found). Inserts highlight affected cells. Nasal turbinates (200x), trachea (200x) and lung tissue (100 and 400x insert) are shown.

### Transmission of MERS-CoV in hDPP4 mice

Next, we investigated the transmission of MERS-CoV within our mouse model via three different routes: fomite, direct contact, and the airborne route.

#### Fomite transmission

MERS-CoV-containing media was pipetted on different objects in the cage, including on two plastic and two metal washers, after which two mice were introduced per cage (*Figure S1*). Upon exposure to 5×10^4^ TCID_50_/cage, 11 out of 18 mice reached endpoint criteria (*Figure 4a*). Brain and lung tissue were harvested from non-survivors, and viral gRNA was detected in brain tissue of eleven mice and lung tissue of seven mice (*Figure 4b*). sgRNA was detected in brain tissue of three mice, and lung tissue of one mouse (*Figure 4c*). Infectious virus was recovered from brain tissue of three mice (*Figure 4d*). The remaining seven mice were euthanized at 28 dpi; six animals were seropositive for MERS-CoV S1 (*Figure 4e*). We then exposed twenty mice to 5×10^6^ TCID_50_/cage, again two mice per cage. Fifteen out of 20 mice were euthanized (*Figure 4a*). Viral gRNA was detected in lung and brain tissue of all the non-survivors (*Figure 4b*). Viral sgRNA was detected in brain tissue of all mice and lung tissue of six mice (*Figure 4c*). Infectious virus was isolated from brain tissue of eight mice (*Figure 4d*). Four out of five surviving animals were found to be seropositive for MERS-CoV S1 protein (*Figure 4e*).

**Figure 4.**
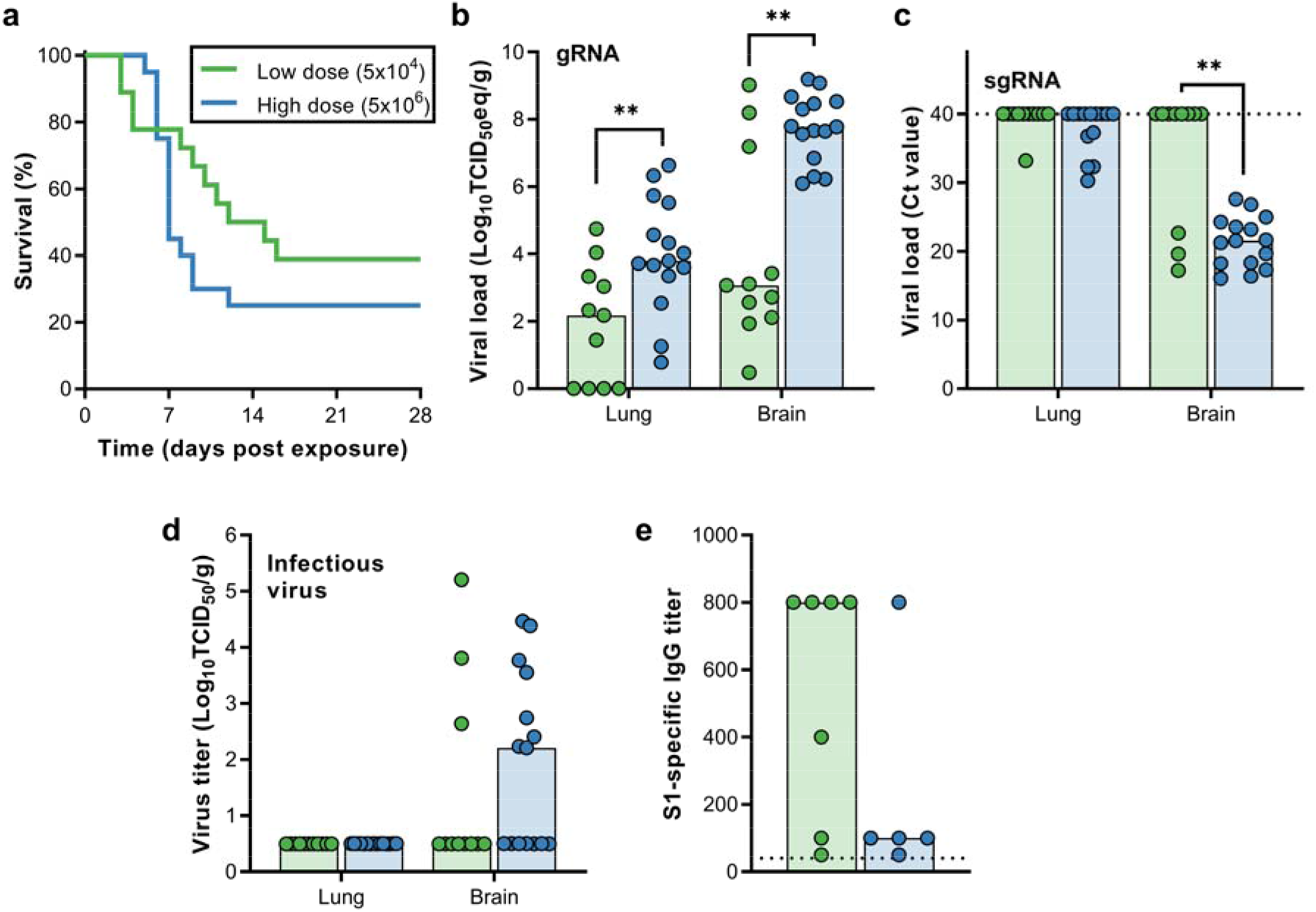
Transmission of MERS-CoV via fomites. hDPP4 mice were exposed to 5 × 10^4^ TCID_50_ MERS-CoV (N=18) or 5 × 10^6^ TCID_50_ MERS-CoV (N=20). (a) Survival curves of hDPP4 mice exposed to fomites containing MERS-CoV. (b-c) Viral gRNA (b) or sgRNA (c) in lung and brain tissue of hDPP4 mice. (d) Infectious MERS-CoV detected in lung and brain tissue of hDPP4 mice. (e) Serology titers in sera of survivors obtained 28 dpe. ELISA assays were performed using MERS-CoV S1 protein. Dotted line = limit of detection. ** = p-value < 0.01.

#### Contact transmission

Twenty mice in two separate experiments were inoculated intranasally with 10^4^ TCID_50_ MERS-CoV, and single-housed. One day later, one sentinel mouse per cage was introduced. No sentinel mice reached endpoint criteria (*Figure 5a*). One mouse seroconverted with a titer of 100 (*Figure 5e*). Subsequently, 38 mice in three separate experiments were inoculated with 10^5^ TCID_50_ MERS-CoV, and sentinel mice were introduced one day later. One sentinel mouse was euthanized five days post-exposure (dpe) (*Figure 5a*). A low amount of viral gRNA was detected in both lung and brain tissue (*Figure 5b*), but no sgNA was detected (*Figure 5c*). Likewise, no infectious virus was recovered from tissue (*Figure 5d*). All 37 remaining sentinels survived exposure. Antibodies against S1 were detected in one surviving animal (*Figure 5e*).

**Figure 5.**
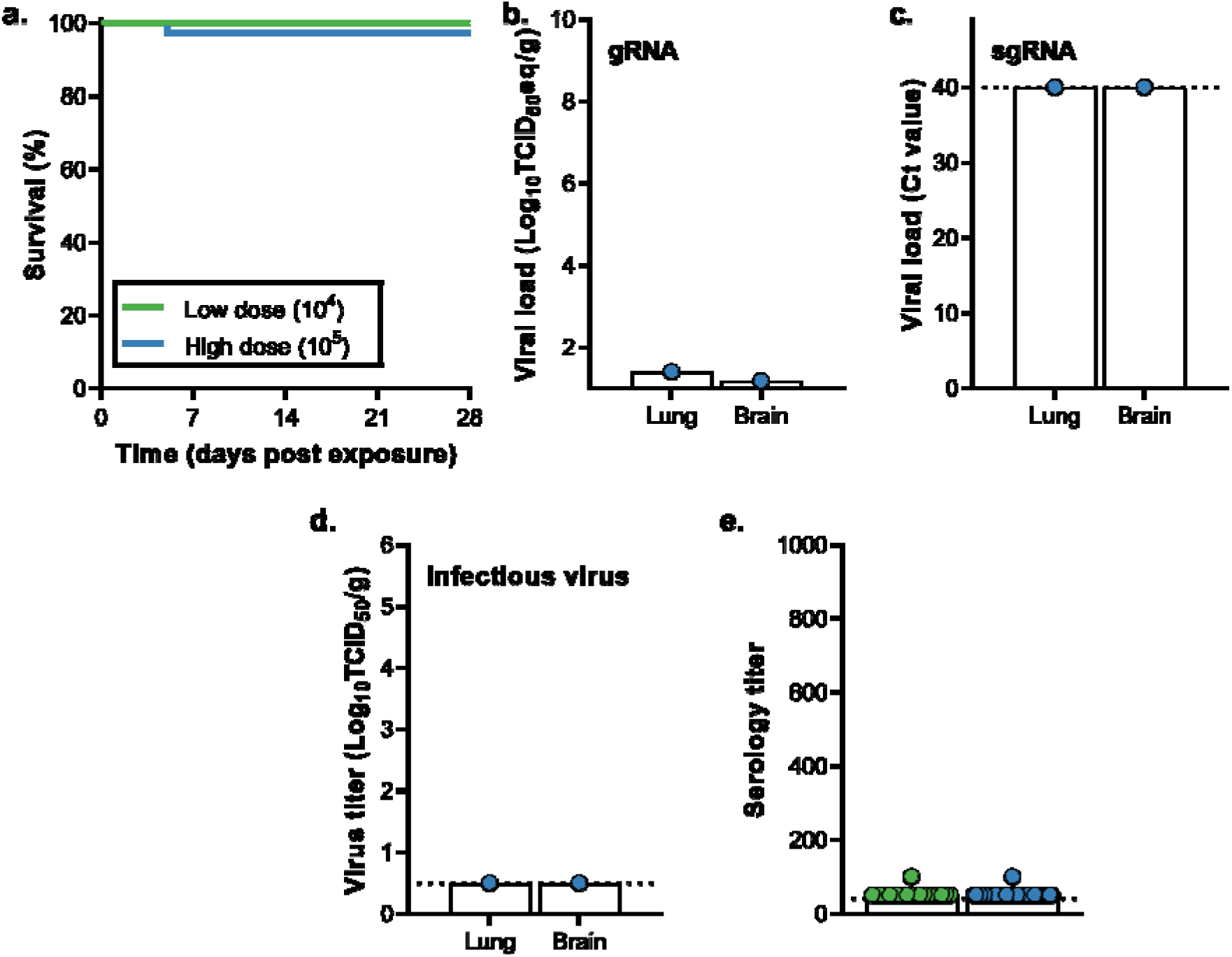
Transmission of MERS-CoV via direct contact. hDPP4 mice were inoculated intranasally with 10^4^ TCID_50_ MERS-CoV (N=20) or 10^5^ TCID_50_ MERS-CoV (N=38). (a) Survival curves of mice directly exposed to donor mice infected with MERS-CoV. (b-c) Viral gRNA (b) or sgRNA (c) in lung and brain tissue of hDPP4 mice. (d) Infectious MERS-CoV detected in lung and brain tissue of hDPP4 mice. (e) Serology titers in sera of survivors obtained 28 dpe. ELISA assays were performed using MERS-CoV S1 protein. Dotted line = limit of detection.

#### Airborne transmission

Five hDPP4 mice were inoculated intranasally with 10^4^ TCID_50_ MERS-CoV. Sentinel mice were introduced in the same cage as the donor animal at one dpi. Animals were separated by a perforated divider, which did not allow direct contact but allowed airflow from the donor to the sentinel animal (*Figure S1*, also described in ^7^). No animals reached endpoint criteria (*Figure 6a*). Three surviving animals were found to be seropositive for S1 at very low titers (*Figure 6e*). In eight different experiments, 54 hDPP4 mice were inoculated with 10^5^ TCID_50_ MERS-CoV. Sentinel animals were introduced one dpi. Four animals reached endpoint criteria (*Figure 6a*). Viral gRNA could be detected in all brain tissues and three out of four lung tissues (*Figure 6b*). Viral sgRNA was detected in brain and lung tissue of two mice, and infectious virus was found in the brain of one mouse (*Figure 6c-d*). All 50 remaining sentinels survived exposure. Seven of the surviving animals were found to be seropositive for S1 (*Figure 5e*).

**Figure 6.**
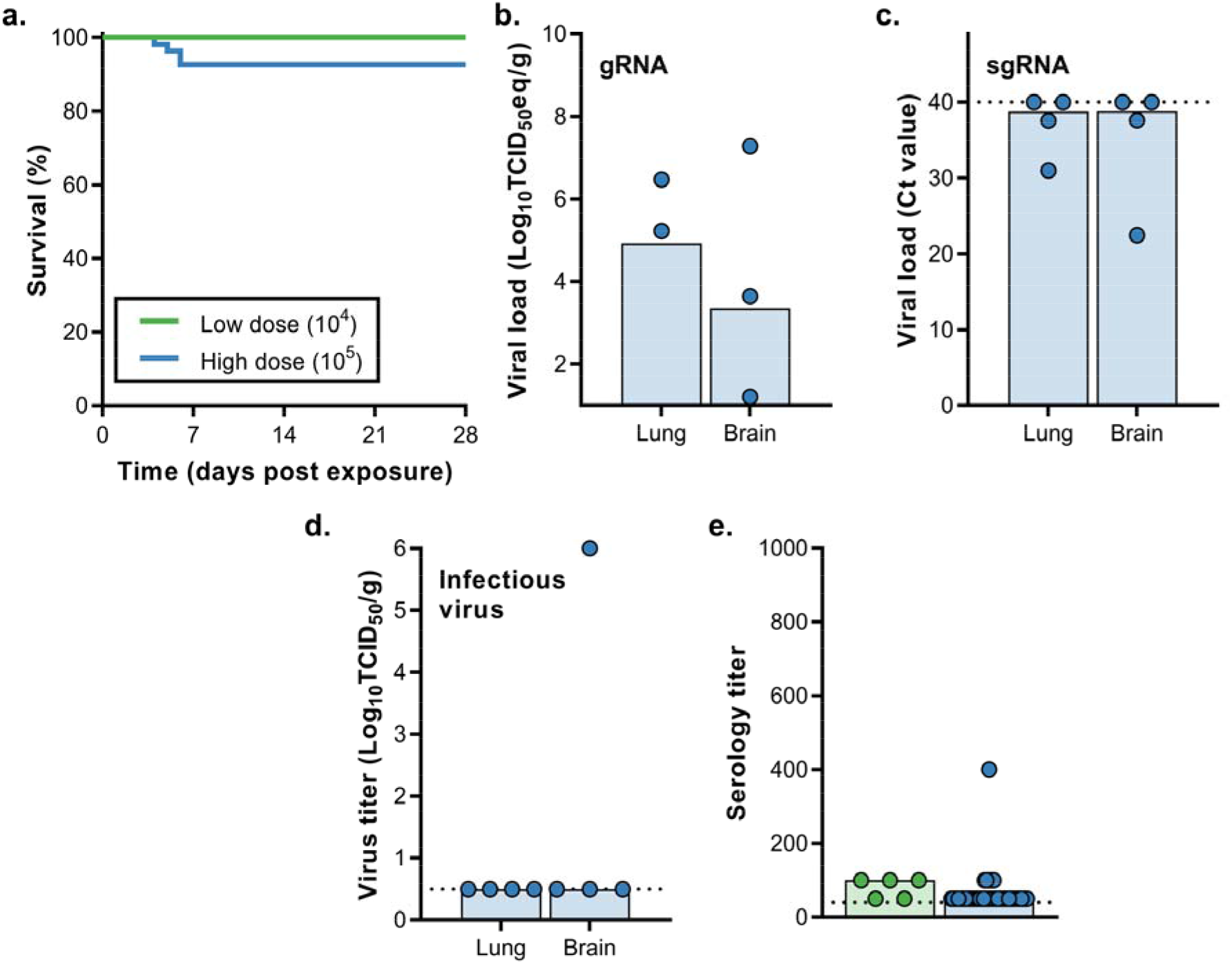
Transmission of MERS-CoV via the airborne route. hDPP4 mice were inoculated intranasally with 10^4^ TCID_50_ MERS-CoV (N=5) or 10^5^ TCID_50_ MERS-CoV (N=54). (a) Survival curves of mice exposed to donor mice infected with MERS-CoV. (b-c) Viral gRNA (b) or sgRNA (c) in lung and brain tissue of hDPP4 mice. (d) Infectious MERS-CoV detected in lung and brain tissue of hDPP4 mice. (e) Serology titers in sera of survivors obtained 28 dpe. ELISA assays were performed using MERS-CoV S1 protein. Dotted line = limit of detection.

## Discussion

The continued circulation of MERS-CoV in the dromedary camel population highlights the need for a better understanding of the transmission potential of MERS-CoV. Like MERS-CoV, SARS-CoV-1 and SARS-CoV-2 are thought to use a combination of different transmission routes between humans, including fomite, direct contact, and airborne transmission. Which one of these routes is most important is difficult to ascertain. However, it is clear that human-to-human transmission of MERS-CoV is restricted compared to SARS-CoV-1 and, in particular, SARS-CoV-2, and is primarily nosocomial ^17^.

Within hospitals, MERS-CoV viral RNA has been detected on various surfaces up to five days after viral RNA was detected in patient samples. In addition, infectious virus was isolated from different hospital surfaces ^30^ and air samples ^31^. Likewise, SARS-CoV-1 and SARS-CoV-2 RNA could be detected in air and surface samples ^32,33^. Experimentally, infectious MERS-CoV at 20°C and 40% relative humidity could be recovered from plastic and steel surfaces for up to 48 hours ^4^, and using a similar setup, SARS-CoV-1 and SARS-CoV-2 likewise retained viability for 48 hours ^5^. Superspreader events have been documented for all three viruses ^34–36^, and in some scenarios, airborne transmission of SARS-CoV-1 and SARS-CoV-2 appears to be the most likely scenario for specific human-to-human transmission clusters ^34,37–39^.

Animal models are crucial for experimental transmission studies, as transmission involves several factors: shedding of virus from an infected host, survival of the virus in aerosols or on surfaces, and infection of the sentinel host. Our data suggest that MERS-CoV can utilize a variety of different transmission routes, although the fomite route was much more efficient than both the direct contact and airborne routes.

In our airborne transmission setup the cage divider prevents direct contact, but allows the movement of larger droplets and aerosols from the donor cage to the sentinel cage. Therefore, we cannot distinguish between transmission events by aerosols (droplets < 100 µm), larger droplets (> 100 µm), or a combination of these two ^40^. An experimental setup exclusively allowing transmission of aerosols as designed for SARS-CoV-2 ^41^ would be able to distinguish between aerosol and droplet transmission.

Our overall data agree with limited human-to-human transmission of MERS-CoV in the general population. Given the tropism of MERS-CoV for the lower respiratory tract and minimal evidence of infection of the upper respiratory tract ^29,42–44^, there is likely little to no natural generation of infectious aerosols in patients. We hypothesize that the propensity of MERS-CoV to transmit relatively efficiently in hospital settings is linked to performing aerosol-generating procedures, such as intubation and bronchoscopy, on infected patients ^45–47^ rather than the natural generation of aerosols containing MERS-CoV from the respiratory tract of the patient. Combined with a potentially more susceptible hospital population ^48^, this could lead to a relatively high nosocomial transmission compared to household transmission.

In this study, we showed MERS-CoV transmission via the fomite and in limited numbers via direct contact and airborne routes. The hDPP4 transmission model will be invaluable in assessing the transmission potential of novel MERS-CoV strains without prior adaptation to the mouse host in light of the continuing virus evolution during human outbreaks and the camel population ^49,50^. In addition, this work will allow the development of effective countermeasures to block human-to-human transmission of MERS-CoV.

## Acknowledgements

The authors would like to thank Laura Tally and colleagues for assistance with mouse breeding, the Rocky Mountain Veterinary branch for assistance with high containment husbandry and cage design, Tina Thomas, Dan Long and Rebecca Rosenke for assistance with pathology and Anita Mora and Ryan Kissinger for assistance with the figures. This research was supported by the Intramural Research Program of the National Institute of Allergy and Infectious Diseases (NIAID), National Institutes of Health (NIH).

## Author contributions

N.v.D. designed the project, performed the experiments, interpreted the data, and wrote the manuscript. T.B., R.J.F., A.O., D.F., R.J.M., and M.L. performed experiments and revised the manuscript. D.S. and G.S. performed experiments, interpreted the data, and revised the manuscript. V.J.M. designed the project, interpreted the data, and wrote the manuscript.

---

**Figure S1.**
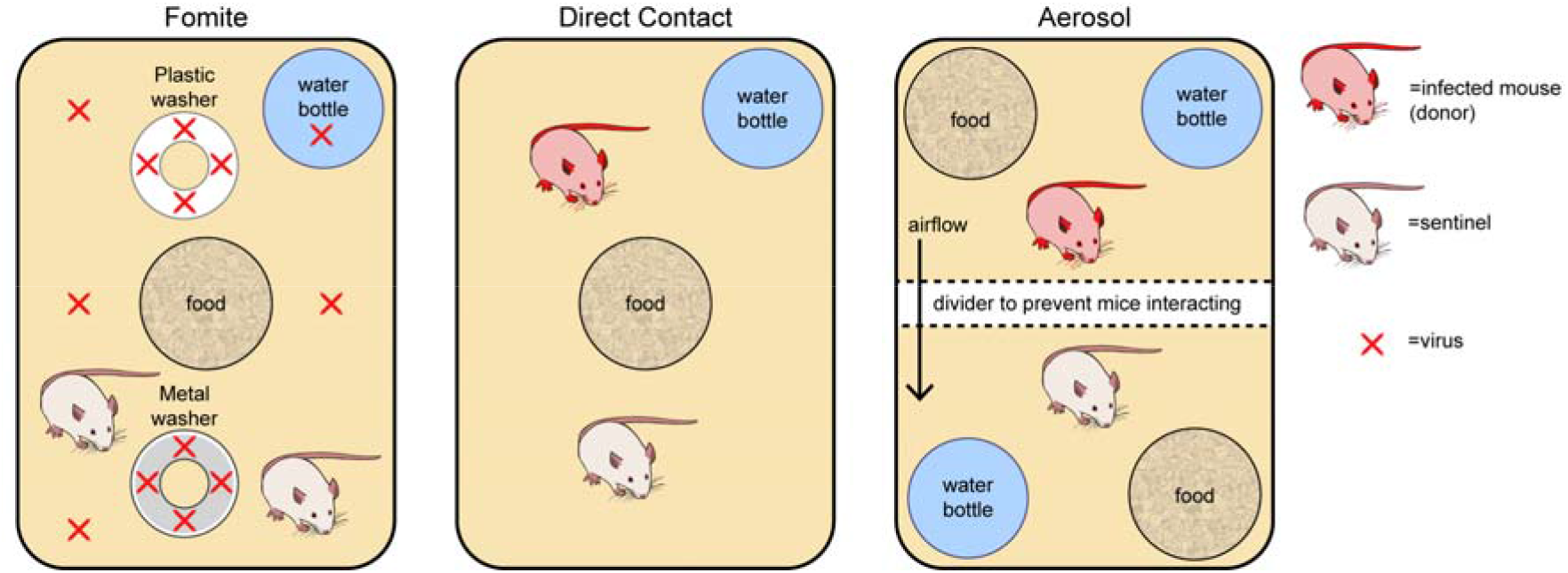
Cage setup during transmission experiments. Depicted are cage setups for fomite, direct contact, and airborne transmission.

